# Chaotic behavior influenced by an environmental parameter in simulated *C. elegans* populations

**DOI:** 10.1101/2025.05.13.653639

**Authors:** Erik Bergstrom, Andrea Scharf

## Abstract

Population dynamics can be chaotic when driven by internal feedback loops, including density-dependent mechanisms, making them predictable only for short periods of time. Since the first discovery of chaos in simple population models, many wild and laboratory populations have been characterized as chaotic. We ask whether a population simulation through a bottom-up modeling approach such as agent-based modeling can display chaotic dynamics. Here, we use wormPOP to simulate *C. elegans* population dynamics with changing culling parameters and calculate the largest Lyapunov Exponents of the simulated populations. The agent-based model wormPOP is a realistic simulation based on a laboratory population system with *C. elegans* and its food source *E. coli*. Our results show that depending on the percentage of culling, populations can exhibit chaotic or periodic behavior. Build-in randomization and the buffering of stress through alternative stress-resistant larva stages in *C. elegans*, complicates the analysis of wormPOP populations, however, the calculated LE for the identified chaotic conditions is robust to changing parameters in our analysis and fits expectations based on the scaling of LE with body size. In conclusion, agent-based models can generate realistic population dynamics that can be chaotic dependent on the chosen parameters.

## Introduction

Chaos has been a relevant concept in ecology since the 1970’s, when Robert May showed that simple population models could display chaotic dynamics^1^. This changed the view that irregular population dynamics are always caused by variable environments or measuring errors. May’s demonstration of chaos in population dynamics lead to the new understanding that traits of individuals in a group can cause irregular fluctuations through internal nonlinear feedback loops such as density dependent behavior^2^. Chaotic systems are generally characterized by sensitive dependence on initial conditions^3^. Small differences between similar states of a chaotic system become exponentially magnified over time at a rate which can be quantified by the largest Lyapunov Exponent (LE) of the system^4^, imposing a fundamental limit on long-term predictability^2,3,5^.

Since chaos was introduced to ecology, mathematical models^6–8^, laboratory populations (for example^9,10^), and time series from wild populations have been used to investigate chaos in populations and ecosystems^11^. Populations and ecosystems that exhibit chaotic behavior have been identified across a wide range of species and situations, from microbial populations in chemostats to communities of larger organisms such as a community of barnacles, mussels, and algae ^9,12–14^. Chaos may be relatively common in natural ecosystems, with one study finding evidence for chaos in >30% of systems^15^. The prevalence of chaos in natural ecosystems makes understanding the impact of chaos on populations and when it occurs a significant concern for conservation, where steady-state management approaches are commonly applied^16^. A variety of methods have been used to identify chaotic behavior, including population models such as the Larval-Pupal-Adult model^13^ and analytical methods such as recurrence quantification analysis^17^ and Rosenstein’s method^18^. Individual-based models have also been used to examine chaos in the context of controlling chaotic population dynamics^19^.

In this study, we analyzed a population time series of *Caenorhabditis elegans* (*C. elegans*) and its food source *Escherichia coli (E. coli)* simulated by wormPOP, an individual-based model^20^. wormPOP is a simulation of laboratory *C. elegans* populations and developed by encoding measured single worm characteristics as rules. *C. elegans* is a free-living nematode that dominantly lives on rotten plant material in the wild and is commonly used as a genetic model organism in biomedical research^21^. The animals are hermaphrodites with a 3-day lifecycle which was conceptualized into an agent-based model with five developmental stages: eggs, larvae, dauer, adult, and parlad (“bag of worms”). After hatching, the animal develops through four larval stages into a mature, egg-laying adult, which will eventually die of old age under optimal conditions. However, when environmental conditions become harsh such as during food scarcity, early larval stages can develop into an alternative diapause stage called dauer, resuming normal development when food is abundant^22^. Under these conditions, adults stop laying eggs leading to the phenomenon where eggs hatch and develop into dauers inside the parental worm before bursting out into the environment^21,23^ – here referred to as parlad. Furthermore, the dauer stage is most likely the dispersal stage in natural *C. elegans* populations^21^. In the laboratory, we hypothesized that dauers persisting through periods of starvation would contribute to stabilizing population dynamics by forming a reservoir of non-feeding dauer larvae that is ready to reenter the development when food is abundant again^20,22^.

*C. elegans* populations are maintained in the laboratory by periodically culling animals and adding bacteria as food^20^. Feeding and culling also occur periodically in wormPOP to mimic the laboratory conditions. During culling, some proportion of the total population is removed from the laboratory population. In wormPOP, this is done by giving each worm a changeable percentage chance to be culled when the culling step occurs. The amount and types of worms periodically removed from the simulation during the culling step can be manipulated, for example to specifically only remove larvae than all life stages, allowing for the impact of culling on different life stages to be examined.

Here, we analyzed simulations with different culling conditions, including all stage culling where all life stages and bacterial food were removed, and dauer & larvae culling where only those dauers and larvae were removed. Time series data from these simulated *C. elegans* populations show a change in dynamics as the proportion of culling increased, with populations with low overall culling showing complicated dynamics and populations with high overall culling following stable periodic oscillations. We applied Rosenstein’s method^18^ to estimate the largest LE, a metric that indicates how rapidly a system diverges, for the simulated populations. Populations at a lower proportion of all stage culling had a higher calculated LE, though all populations had a positive LE by this method of analysis. Testing whether the LE decreased as the embedding dimension, a parameter for Rosenstein’s method, increased indicated that the group of conditions with lower calculated LE values were likely falsely identified as positive due to the presence of noise in the time series. The calculated LE values for the chaotic conditions where the LE did not decrease as the embedding dimension were similar to the value that could be expected based on the correlation between the body size of a species and the LE of populations of that species^24^.

## Materials and Methods

### Simulation Methods

Population time series were generated using wormPOP version 1.0^20^ with varying culling conditions, using the standard parameters for wild-type worms as described in Scharf et al., 2022^20^. The populations were culled and fed every 24 hours. The proportion of culling was incremented by 5% starting at 0% to a maximum 40% for overall culling. For dauer and larval culling, the culling proportions used were 25%, 50%, 75%, 80%, and 84.5%. The dauer and larval culling conditions close to 85% were chosen since previous research^20^ had found a tipping point at 85% culling where worms switched from dying of starvation to dying of old age. Feeding stayed constant with 10 mg bacteria every 24 hours. The number of timesteps was increased to 20,000, which is equivalent to 2,500 days, to collect an adequate number of data points for analysis.

### Calculation of Lyapunov Exponents

LE values were determined using the method of Rosenstein et al ^18^ using the number of worms from every eighth time point of the time series, which is equivalent to one time point per day. Every eighth timepoint was used to avoid complications in data analysis due to the feeding times in the simulation occurring every 8 timesteps, the equivalent of once per 24 hours. The lag was chosen by using autocorrelation and the embedding dimension was chosen the method of false nearest neighbors, as recommended by Marwan in 2011^25^. A Theiler window of T = 6 was used based on space-time separation plots^26^ using the subset time series from populations at each condition. A Kruskal-Willis test was used to test whether the calculated LE values were the same among all conditions, then a Dunn pairwise test was used to conduct comparisons among conditions.

### Test for positive LE values caused by stochastic variation

Methods to directly estimate the LE from a time series often produce positive LE values when noise is included in a time series^27^. Since several events in wormPOP including culling, starving of larvae, and exiting/entering dauer are at least partially stochastic, it was necessary to test whether the determined LE values here were due to chaotic dynamics or noise. LE values were calculated for each embedding dimension from 10 to 20 for each time series, then a linear regression was used to determine the change in LE as the embedding dimension was increased. Assuming the initial embedding dimension is sufficient, time series dominated by stochasticity are expected to show a clear decrease in LE value as the embedding dimension is increased whereas time series dominated by chaotic behavior are expected to show no change in LE as the embedding dimension is increased(10). A Dunn pairwise comparison test with holm correction was used to test whether the identified slopes were significantly different from zero.

## Results and Discussion

### *C. elegans* populations show different patterns and levels of synchrony between simulations

To compare how simulated *C. elegans* populations differ when started with the same parameters, we simulated ten populations using identical parameters. Furthermore, we were interested in whether the heterogeneity between populations changes with variations in the culling parameters, and therefore selected six different culling conditions: 0%, 25%, and 30% culling all stages and 25%, 75%, as well 84.5% only dauer & larvae culling (Fig. 1). All populations were initiated with a founding population of 1000 animals and exhibit exponential growth to a first peak. The time to first and height of this first peak varied from 10 days at 0% culling to a maximum of 93 days at 84.5% larval and dauer culling. Populations showed little variance in the time and height of the first peak (Fig. 1A-E) except for the populations with 84.5% larvae and dauer culling (Fig. 1F). Here, the timing of the first peak varied between 81 and 93 days. After the first peak, these worm populations mostly stayed stable with only some fluctuations around 50,000 worms, whereas populations with all other culling parameters show much higher fluctuations. Specifically, populations with a culling rate below 25% exhibited varied patterns after the first population peak (Fig 1 A,B,D). Different simulations with these conditions did not show the same populations fluctuations and populations desynchronized in the first 25 days. In contrast, populations with more than 25% all stage culling as well as more than 50% dauer & larval culling exhibited somewhat periodic population cycles after the initial population peak (Fig. 1C,E). Populations with 75% dauer & larval culling were unique in having multiple periodic cycles and continued to remain in the same cycles after the initial population peak for the entire length of the time series collected. Previous studies on flour beetles demonstrated that laboratory populations operated within a chaotic attractor where they unpredictably switched between multiple periodic cycles^13^. In contrast, simulated worm populations with 75% larval & dauer culling exhibited one of two cycles after the initial population peak and continued in that cycle for the remainder of the simulated time series. In summary, simulated *C. elegans* populations exhibited different patterns of population behavior and synchrony between conditions.

**Figure 1.**
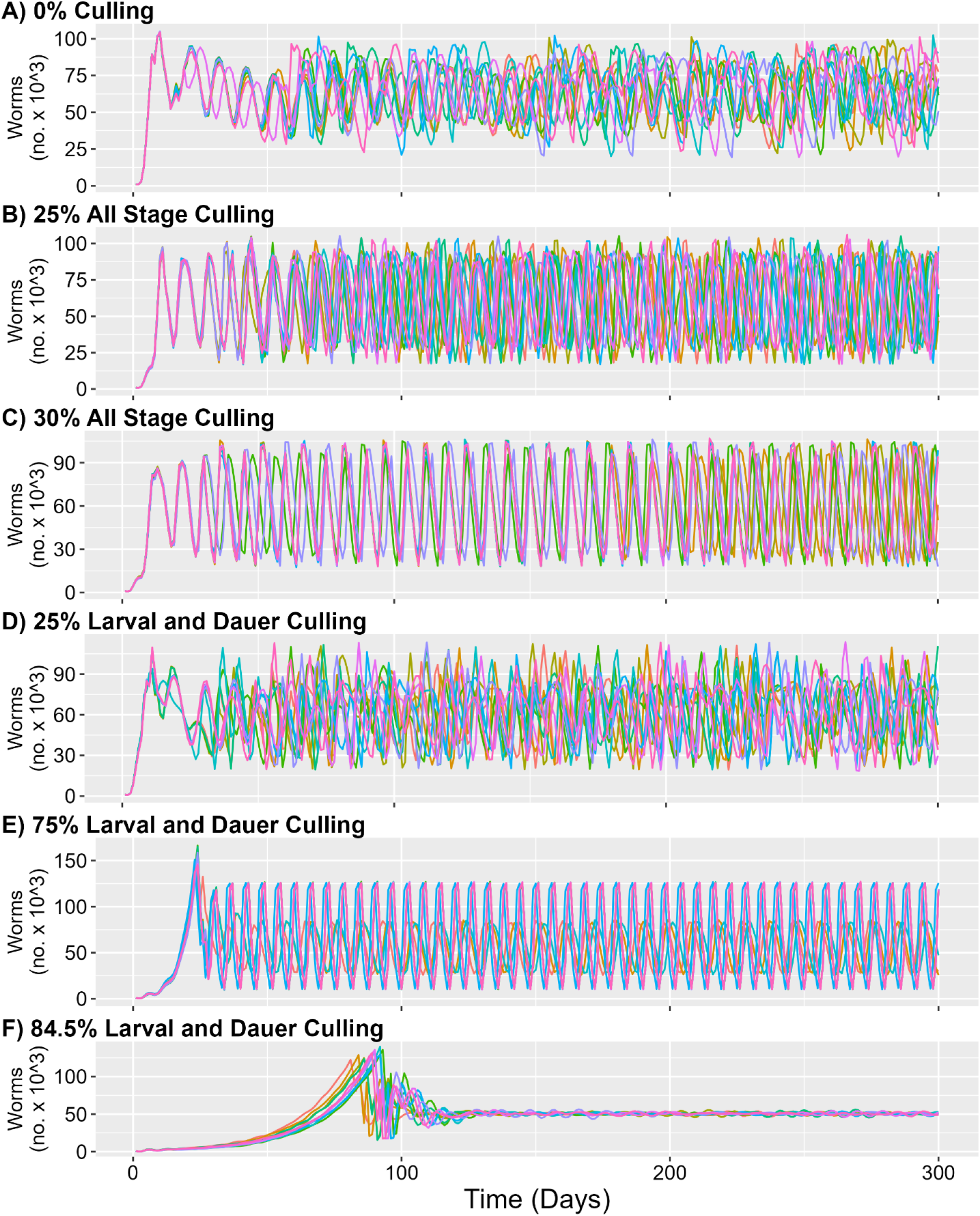
Simulated *C. elegans* population dynamics with different culling conditions. The graphs show the number of worms over the first 300 days of ten independent computational simulations with (A) 0% all stage culling, (B) 25% all stage culling, (C) 30% all stage culling, (D) 25% larval and dauer culling, (E) 75% dauer and larval culling, and (F) 84.5% larvae and dauer culling. The different colors represent the different individual runs. Note that with some conditions the individual simulations are very similar and the colors overlap.

### Variation in culling parameters changes the population behavior from chaotic to more stable population dynamics

Next, we calculated the largest LE for all conditions using the method described in Rosenstein^18^ to determine whether the observed population dynamics were chaotic (Figure 2). The calculated LE values for 0-25% overall culling and 25% dauer & larval culling were significantly higher than calculated LEs for the conditions with higher culling, matching the conditions where worm populations displayed periodic behavior after the first population peak. The spectrum of population behavior observed here is comparable to the simple population models Robert May examined^1^. In his work, population models transitioned from chaotic dynamics to periodic cycles, and then to stable populations at carrying capacity as the rate of growth fell below certain values. In this study, as the percentage of culling is increased, the simulated *C. elegans* populations appear to transition to more stable periodic dynamics and eventually reach a nearly steady equilibrium. Increased culling reduces the rate at which growing *C. elegans* populations can expand, so a similar effect may be causing the changes in population dynamics simulated with wormPOP.

**Figure 2.**
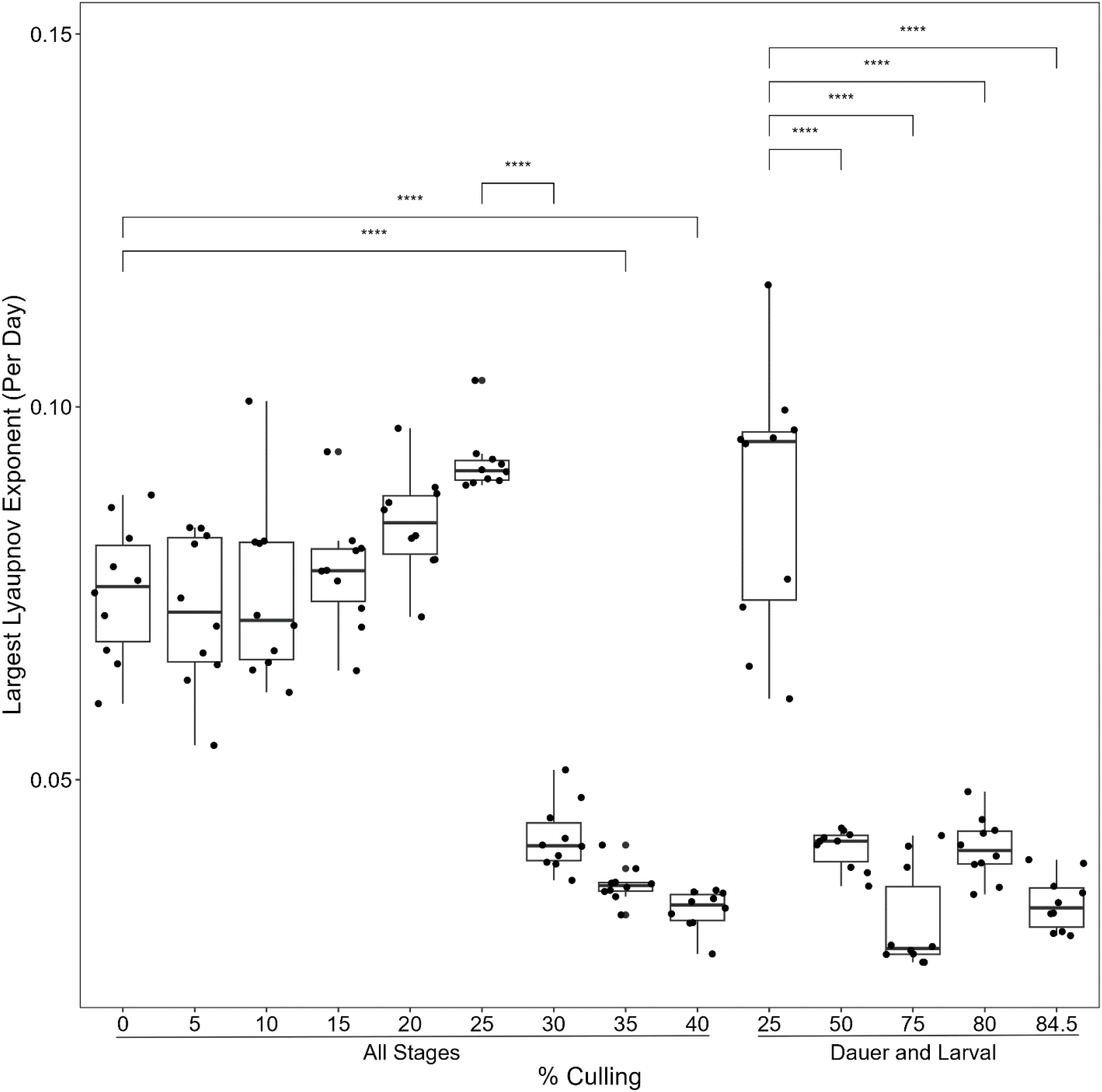
Estimated Lyapunov Exponents of simulated population dynamics with different culling conditions. Calculated LE values for simulated *C. elegans* populations with all stage culling conditions and with only larvae & dauer culling conditions. LE values for simulated *C. elegans* populations with 0%, 5%, 10%, 15%, 20%, and 25% all stage culling, and 25% larval & dauer culling conditions were significantly different from 35% and 40% overall culling conditions as well as 50%, 75%, 80%, and 84.5% larval and dauer culling conditions (Dunn multiple comparisons test with holm correction, p < 0.01).

### Can stochasticity influence the calculated Lyapunov Exponents?

Several conditions, all stage culling above 30% and dauer & larval culling above 25% produced time series that visually appeared to be periodic but showed a positive LE (Fig. 1 and Fig.2). Since Rosenstein’s method can produce positive LE values for stochastic time series^18^ and since wormPOP has stochastic processes^28^, we decided to determine whether the LE values identified here resulted from chaotic dynamics or influenced by noise. When Rosenstein’s method^18^ produces a positive exponent for a stochastic time series, the calculated LE decreases as the embedding dimension, a parameter of the time-delay embedding that is used to prepare the data, is increased. Therefore, one way to test whether the calculated LE values were impacted by noise is to examine how the LE changes as the embedding dimension is increased. Therefore, LE values were calculated as the embedding dimension was increased from 10 to 20 for all culling conditions, and a linear regression was performed to quantify how the LE changed as embedding dimension increases. The slope of the calculated regression line of LE vs. embedding dimension was significantly less than zero for 10%, 25%, 30%, 35%, 40% all stage culling, as well as 50%, 75% and 84.5% dauer & larvae culling (Fig. 3). All other conditions were not significantly different from zero. Time series dominated by chaotic behavior were expected to show no change in LE as embedding dimension is increased^18^. Overall, the results shown in Fig. 4 support therefore the hypothesis that the positive LE values identified for conditions above 25% all stage culling and 25% dauer & larvae culling are due to stochasticity, not actual chaos. Almost all conditions expected to be periodic based on visual inspection of the time series were identified as having a slope that was significantly less than zero, indicating that the positive LE was due to stochasticity in the data. (Fig. 3).

**Figure 3.**
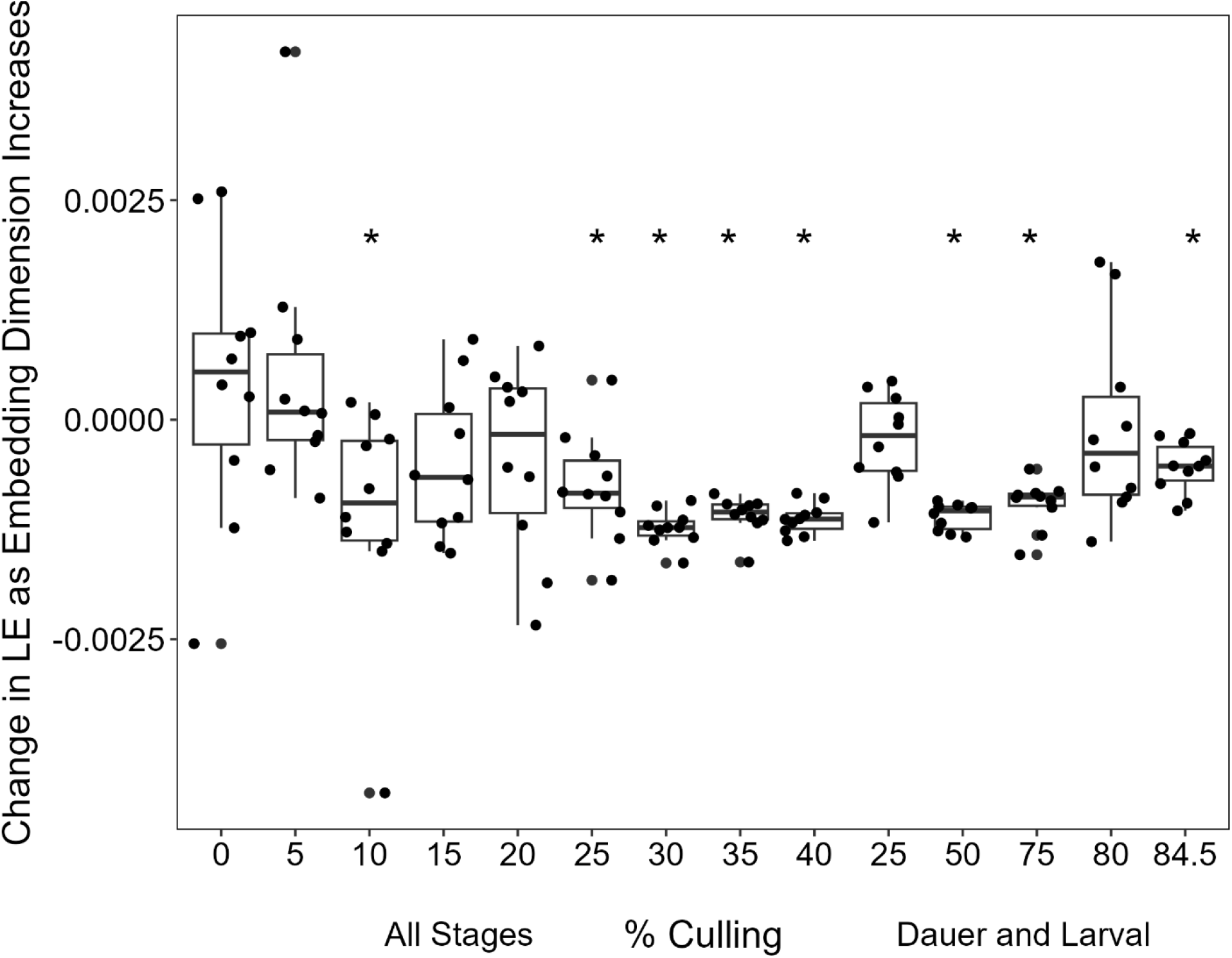
Change in LE as the embedding dimension is increased. Average changes in LE as the embedding dimension was calculated for simulated population dynamics with all stage culling conditions and with only dauer & larvae culling conditions by using linear regression to fit a best fit line to a graph of LE values versus embedding dimension for each time series. The average slope of each culling condition was compared to zero (Wilcoxon-signed rank test, adjusted using the Holm method, p < 0.05). 10%, 25%, 30%, 35%, and 40% all stage culling and 50%, 75%, and 84.5% dauer and larval culling had slopes.

**Figure 4.**
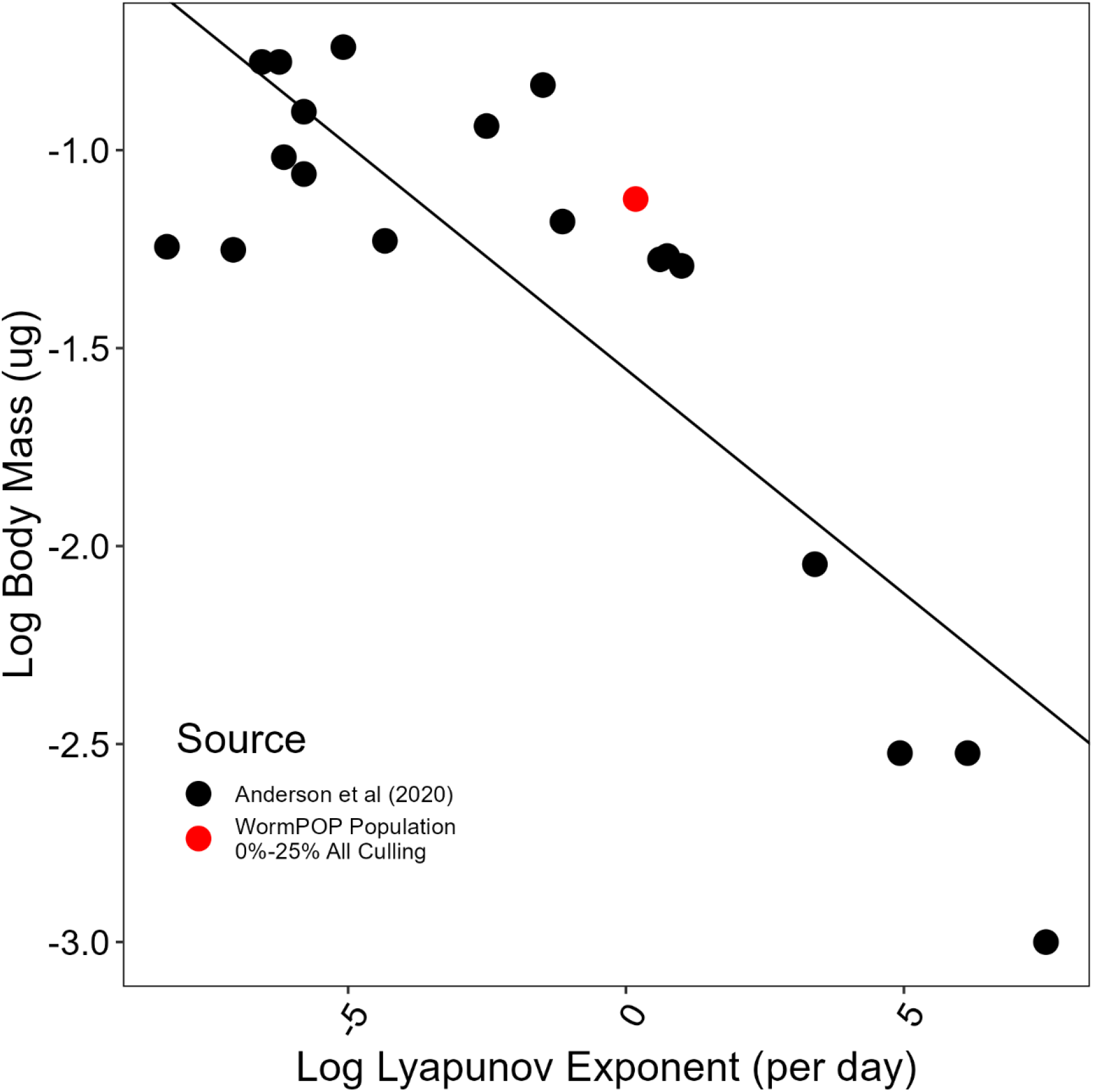
The determined LE for simulated *C. elegans* populations scale with body size. Comparison of calculated LE value versus body mass according to Anderson et al., 2020. The value used for *C. elegans* mass is 1.4 micrograms, which was determined by calculating the average mass of an adult worm in the wormPOP simulation after the initial population peak. The LE value plotted for the wormPOP population corresponds is the average LE value of the 0% all stage culling condition. The data points from Anderson et al. (2020) are the LE values and body mass reported in table one^29^.

Additionally, the slopes of the LE vs. embedding dimension regression line were not significantly different from zero for conditions that appeared to display non-periodic behavior. The slopes at 10% and 25% culling were significantly less than zero, indicating that stochastic events in wormPOP are influencing the LE at these culling conditions. However, the calculated LE values for these conditions are not significantly different from the LE values for conditions that were identified as chaotic. Additionally, the behavior of worm populations at 10% and 25% all stage culling also was also far more dynamic than the behavior of populations at above 25% all stage culling and 25% larval and dauer culling, and the LE of these conditions decreased less uniformly as the embedding dimension increased. These observations indicate that stochastic events influence the LE calculated for 10% and 25% all stage culling, but that these conditions are different from the other conditions with a slope significantly less than zero. One possible explanation is that populations at 10% and 25% all stage culling display chaotic behavior but are also influenced by stochastic events encoded in wormPOP. The slope for the 80% dauer and larval culling condition was unexpectedly not significantly different from zero, despite populations in this condition appearing to display clearly periodic behavior. Some possible interpretations of this result are that populations behave chaotically in this condition despite following mostly periodic cycles, that the initial embedding parameters for this condition were not sufficient to accurately determine the LE. Utilizing an alternative method for detecting chaos may help resolve what type of behavior is occurring under this condition.

It is notable that the calculated LE value for populations with 0% all stage culling is not significantly different than the LE of the populations with other all stage culling conditions below 30% (Fig. 2). Systems with added noise diverge at faster rates, since random events produce differences that become magnified^11^. If the increased LE values for conditions with 5-25% overall culling were due to stochastic variation being introduced by culling, 0% all stage culling would be expected to have a lower LE.

### Simulated populations with 0-25% culling fit into known ecological patterns

How do the ecological patterns found in populations simulated with wormPOP fit into previously described patterns? It has been shown that Lyapunov exponents of populations scale with body size^15,24^. The data from Anderson and colleagues includes LE values derived from direct estimation using laboratory population time series, as well as population-level modeling^24^ (Fig. 4). We compared the here calculated LEs of populations with culling conditions identified as chaotic to the data from Anderson et al., 2020 to see whether our reported LE fit into the reported scaling trend. As shown in Fig. 4, the LE is generally consistent with the reported trend of LEs decreasing as body size increased, which is an indication that the method used in this paper of using an individual-based simulation to produce time series for direct estimation of the LE could be useful for future research projects. One benefit of utilizing an individual-based simulation is that it allows for individual details, such as body mass, to be accurately tracked during an experiment. Another application of wormPOP or similar simulations would be to investigate how individual traits, such as life history, affect the occurrence of chaos in populations, since these traits can be altered much more easily in individual-based simulations. The need for comprehensive data on the behavior of individual organisms is a limiting factor for this method, unless general models such as the dynamic energy budget^30^ are applied.

### The complex lifecycle of *C. elegans* complicates comparisons to simple population models

Thus, our results indicate that changing the levels of culling induced significant changes in the largest LE for both all stage culling and dauer & larvae culling. Previous studies discovered that manipulating environmental parameters can influence the behavior of a system. For example, Benincá and colleagues^9^ found that changes in the dilution rate at which material was added to and removed from a chemostat containing a plankton community could result in the community displaying both periodic and chaotic dynamics^31^. The dilution rate parameter of a chemostats can be compared to the culling parameter of the here simulated *C. elegans* populations, as both control the rate at which organisms leave the system. Unlike Benincá et al’s population experiments, however, all culling conditions in this study produced a positive LE. Interestingly, using other time series from wormPOP instead of the number of worms generated similar results, with conditions below 30% all stage culling along with 25% dauer and larval culling having a higher LE than the other conditions.

In summary, our results for simulated *C. elegans* populations are very similar to those found by Robert May’s initial mathematical models^1^, where a simple model of population growth was found to exhibit chaotic behavior for a sufficiently high growth rate. A higher proportion of culling in wormPOP simulations effectively reduces the maximum growth rate of the *C. elegans* populations, which could be one reason for the similarity in behavior. However, this comparison is complicated by the additional complexity of population behavior in wormPOP, especially the presence of stress-induced dauers. Dauers regularly occur in simulated populations and they have the potential to stabilize population dynamics. They can survive long periods of starvation before exiting and resuming development into a reproductive adult and form therefore a reservoir that buffers against strong population drops and extinction^20,23^. However, the presence of dauers clearly did not prevent the populations from displaying chaotic dynamics with culling rates under 10% all stage culling. It is possible that the feeding schedule in WormPOP mitigated the dauers impact on population dynamics by preventing prolonged periods of starvation. Future experiments with changes in the feeding schedule such more time between feeding could test whether this is the case.

## Conclusion

Our work provides evidence for chaos in simulated *C. elegans* populations and evidence for chaos originating from the emergent dynamics of an individual-based model. Most models used in ecology are either mathematical models or population-based models that predict population growth and behavior without involving individual traits. This individual-based model generates emergent population dynamics predicted by some ecological models, helping to bridge the gap between theory and laboratory experiments.

## Acknowledgements

We thank Josh Mitteldorf for wormPOP support and discussion, Kerry Kornfeld for comments and discussion and the wormPOP team members, Clare Koerkenmeier, Molly Ripper, and Phuong Tran for preliminary data. This work was supported by the Missouri S&T OURE Fellows Program.

